# A Logic-Based Dynamic Modeling Approach to Explicate the Evolution of the Central Dogma of Molecular Biology

**DOI:** 10.1101/103127

**Authors:** Mohieddin Jafari, Naser Ansari-Pour, Sadegh Azimzadeh, Mehdi Mirzaie

## Abstract

It is nearly half a century past the age of the introduction of the Central Dogma (CD) of molecular biology. This biological axiom has been developed and currently appears to be all the more complex. In this study, we modified CD by adding further species to the CD information flow and mathematically expressed CD within a dynamic framework by using Boolean network based on its present-day and 1965 editions. We show that the enhancement of the Dogma not only now entails a higher level of complexity, but it also shows a higher level of robustness, thus far more consistent with the nature of biological systems. Using this mathematical modeling approach, we put forward a logic-based expression of our conceptual view of molecular biology. Finally, we show that such biological concepts can be converted into dynamic mathematical models using a logic-based approach and thus may be useful as a framework for improving static conceptual models in biology.

## Introduction

In 1965, the pioneering work of Jacob and Monod showed that DNA is transcribed to RNA and further translated into protein, and that the rate of transcription is controlled by a feedback loop in which protein regulates the activity of the transcriptional complex [1]. This vignette was also meticulously illustrated in the three kingdoms of life by Francis Crick [2] where he formulated an information transfer law in biological systems, namely the Central Dogma (CD) of molecular biology [3].

Analogous to the “gene” definition, which has been updated continuously in the past century and a half [4], this fundamental biological law has also been modified [3, 5, 6] since the flow of biological information is understood to be much more complex than previously thought. It has been discovered that DNA itself is not static and genes may be permanently silenced based on the needs governed by cellular circumstances [7]. Proteins may also transfer information in the alternate direction by silencing genes through epigenetic mechanisms. In addition, post-translational chemical modifications may switch a protein to active and inactive states, or may result in proteasomal degradation, entailing ubiquitin tagging followed by entering cellular recycling bins, i.e. proteasomes [3]. Moreover, genes are transcribed in the form of distinct splice variants, where exons are omitted or alternative combinations of exons are utilized, leading to a diverse set of gene products with functional differences. The latest species discovered in the field is microRNA (miRNA) which has led to further elucidation of the regulatory process of transcription in which information flow is blocked either by degrading the transcribed mRNA or by facilitating gene silencing at the DNA level [8-11] (Fig. 1A).

**Fig. 1.**
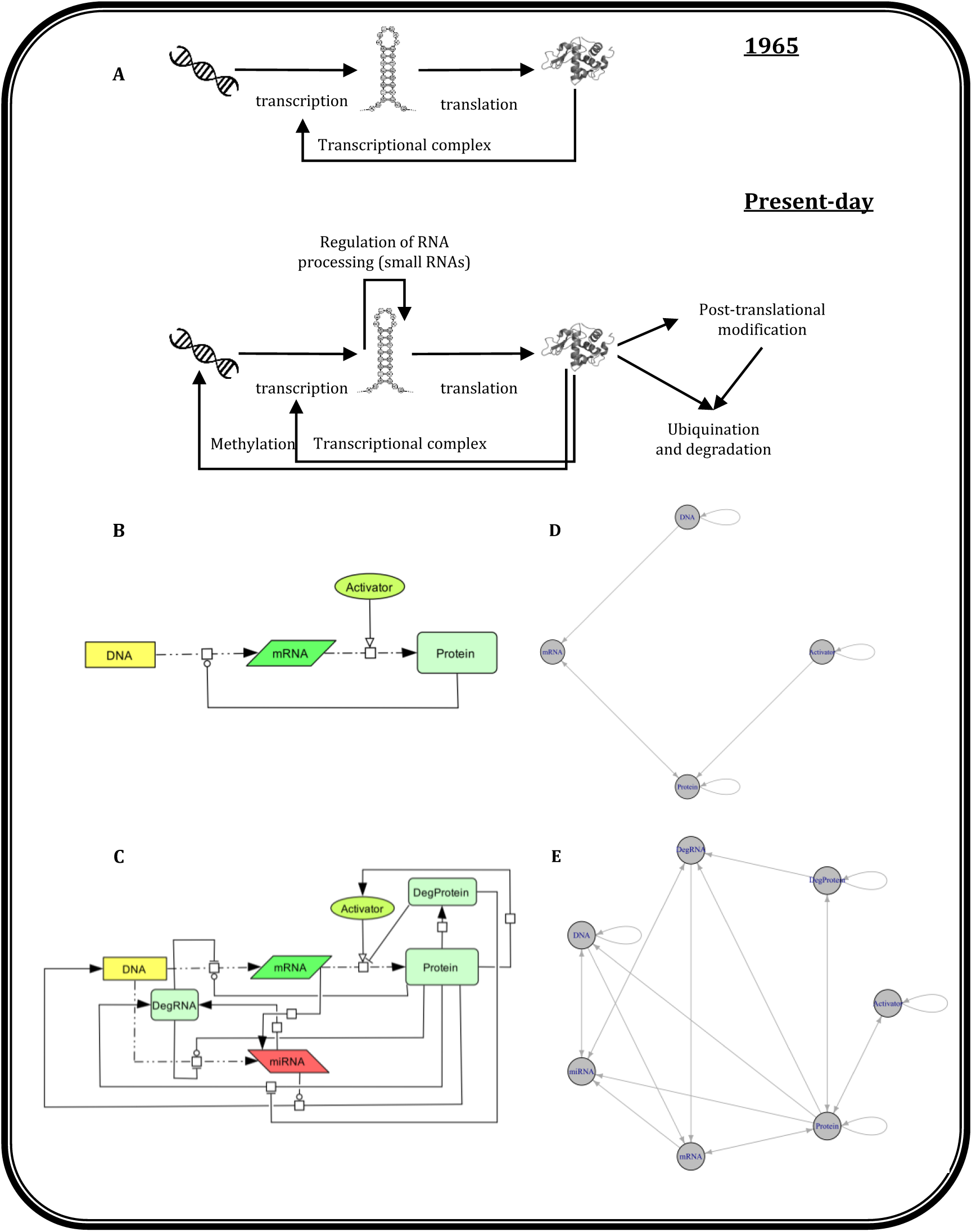
(A) The 1965 and the present-day Central Dogma editions in molecular biology (B) The initial four-node Central Dogma wiring diagram formed by DNA, mRNA, Protein and Activator. (C) The present-day seven-node Central Dogma wiring diagram formed by DNA, mRNA, Protein, Activator, miRNA, DegProtein and DegRNA. The directed edge “—o” represents a functional relationship with both activatory and inhibitory effects (D) Network representation of Boolean rules corresponding to the simple 1965 model. (E) Network representation of Boolean rules corresponding to the present-day updated model. For simplicity, the first version of each model is shown.

The objective of this study was to compare these two different abstractions of information flow reported in 1965 and the present-day revised edition using a dynamic modeling approach. For this, we formulated the following questions. What is to be expected when this conceptual wiring diagram changes in a cell over time? Further, which of the known dynamic behaviors is detectable and which ones are of predictive value? On the basis of the availability of biological information and the nature of the problem at hand, different types of dynamic modeling such as ordinary differential equation (ODE), Wilson-Cowan and fuzzy logic models are used to model a biological question [12-14]. Given the fact that the models presented in this study demonstrated a qualitative problem of certain general concepts in molecular biology, we thus decided to apply discrete logical modeling with two states (i.e. Boolean modeling) to fathom potential dynamic behaviors. Even though Boolean modeling is the simplest species among dynamic modeling methods [15], Robeva and Murrugarra have previously illustrated that it can successfully exhibit complex dynamic behaviors such as bistability [16]. This capability mainly originates from the strength of the step function as a good approximation of the sigmoidal kinetic function of molecular bindings (e.g. enzyme-substrate interaction) and that thresholds exist in most biological processes [17-19]. In line with the abovementioned questions, we aimed to ascertain the potential of logic-based modeling to map out the perceptual design of scientific notions.

## Materials and Methods

The following is the description of the CD models, used in this study, in which, all possible behaviors of single genes and encoded gene products are simulated without considering the combinatorial behaviors of multiple genes for simplicity.

### Boolean dynamic models

The initial representative model of CD containing four components, namely DNA, mRNA, Protein and Activator was reconstructed according to the original studies on CD in molecular biology [1, 2] (Fig. 1B, 1D). Given the activatory or inhibitory effect of Protein on mRNA transcription, two possible versions of this model were constructed (Table 1). In this model, the long half-life components (i.e. DNA, Protein and Activator) are distinguished by a self-loop in Boolean rules (note that self-loops are not shown in Fig. 1B, 1C). It is crucial to emphasize that the names used in the model do not fully comply with the meanings directly inferred in molecular biology. For instance, as described in the second column of Table 1, DNA refers to any gene which can be transcribed to coding RNA. Such definitions are according to the concepts put forth by Francis Crick [2] when splicing, post-transcriptional and translational modifications were still nebulous. Moreover, in this context, the label “Protein” denotes the active and functional protein and not simply any peptide sequence translated by ribosomes.

**Table 1:** Boolean rules governing the state of the 4-node network of the 1965 proposed Central Dogma depicted in Fig. 1B derived from [1, 2]. The symbols “&”, “|” and “!” represent “AND”, “OR” and “NOT” operators.

|  |  | *Logical rule* |  |
| --- | --- | --- | --- |
| *Node* | *Definition* | V1: Activatory gene expression | V2: Inhibitory gene expression |
| *Activator* | The chemical molecules which facilitate active protein production during or after translation | Activator |  |
| *DNA* | The genes transcribed to coding RNA | DNA |  |
| *mRNA* | The coding RNA | DNA & Protein | DNA & !Protein |
| *Protein* | The functional active protein | (mRNA & Activator) \| Protein |  |

In the current edition, CD is more complex given the addition of new components and functional relationships (see Fig. 1C, 1E and Table 2) where the main difference pertains to the concept of “turnover” of the dynamic molecules included in the present-day model of CD. The DegRNA and DegProtein components are thus used to represent the degradation of mRNA and proteins (i.e. DegProtein, DegRNA and Protein) respectively. Yet another revision is the inclusion of miRNA molecules in the model as a representative of all RNA species which negatively affect protein production by regulating mRNA half-life. Regulatory effects of miRNAs on target mRNAs are not exclusively at the post-transcriptional level (miRNA-mRNA) and also show their effects in the form of miRNA-DNA interactions [9-11, 20]. The definition of DNA and Activator has also changed after the emergence of epigenetics and discovery of the post-translational modification phenomenon. The “DNA” label is, therefore, used in this model to explain the non-silenced genes recognized by RNA polymerase and the “Activator” is a molecule that connects to “Protein” in a feedback loop to delineate the role of the latter in the “Activator” turnover. Specifically, the exogenous and endogenous activators are usually affected by proteins such as enzymes, transporters and channels. The rules of this model are derived from the vignette presented in [1-3, 7-11, 20, 21]. Based on the three functional relationships (Protein-mRNA, Protein-miRNA and miRNA-DNA) with each able to take opposite signs (i.e. activation and inhibition) generates eight different versions of present-day CD (Table 2, Supplementary file 1).

**Table 2:** Boolean rules governing the state of the 7-node network of the present-day Central Dogma depicted in Fig. 1C derived from [1-3, 7-11, 20, 21].

|  |  | Logical rule |  |  |  |  |  |  |  |
| --- | --- | --- | --- | --- | --- | --- | --- | --- | --- |
| Node | Definition | V1:<br>Activatory | V2: mRNA<br>expression<br>inhibition | V3:<br>miRNA<br>expression<br>Inhibition | V4: Gene<br>silencing | V5: Gene<br>silencing<br>& mRNA<br>expression<br>inhibition | V6: Gene<br>silencing<br>& miRNA<br>expression<br>inhibition | V7: miRNA<br>& mRNA<br>expression<br>inhibition | V8:<br>Inhibitory |
| Activator | The chemical molecules or residue which facilitate active protein generation before or after translation | Protein \| Activator |  |  |  |  |  |  |  |
| DegProtein | All biochemical modifications that are involved in negative regulation of protein amount | Protein & !DegProtein |  |  |  |  |  |  |  |
| DegRNA | The protein complex which degrades RNA species | miRNA & Protein & !DegProtein |  |  |  |  |  |  |  |
| DNA | The non-silenced genes which is detectable by RNA polymerase | (miRNA & Protein) \| DNA |  |  | (!miRNA & Protein) \| DNA |  |  | (miRNA & Protein) \| DNA | (!miRNA & Protein) \| DNA |
| miRNA | A RNA species which is involved in regulation of gene silencing (for simplification, all other RNA species which has negative effect on protein production is shown with the same label) | (DNA & Protein \| mRNA) & !DegRNA |  | (DNA & !Protein \| mRNA) & !DegRNA | (DNA & Protein \| mRNA) & !DegRNA | (DNA & !Protein \| mRNA) & !DegRNA | (DNA & Protein \| mRNA) & !DegRNA | (DNA & !Protein \| mRNA) & !DegRNA |  |
| mRNA | The mature messenger RNA that is ready for translation (for simplification, all other RNA species which has positive effect on protein production is shown with the same label) | DNA & Protein & !DegRNA | DNA & !Protein & !DegRNA | DNA & Protein & !DegRNA | DNA & Protein & !DegRNA |  | DNA & !Protein & !DegRNA |  |  |
| Protein | The functional active protein | (mRNA & Activator \| Protein) & !DegProtein |  |  |  |  |  |  |  |

### Model simulation and identification of attractors

To identify the synchronous and asynchronous attractors, we used the R package “BoolNet”[22] which conducts an exhaustive search in identifying attractors. The difference between these attractors is based on the updating procedure of components in each time step of the simulation. Therefore, to check the dynamic behavior of components, which are updated either at the same time or at different points of time, we undertook both synchronous and asynchronous updating simulations [15, 19, 23]. The synchronous analysis was separately undertaken on all the ten proposed versions whereas the asynchronous analysis was just done on the eight present-day model versions. In the synchronous simulation, the state-transition graphs were analyzed and the attractors, proportionate with the size of the basin of attraction, were identified. For the asynchronous simulation, the complex attractors were depicted in which betweenness and closeness centralities of each state (node) were illustrated by node size and color. Network visualization and centrality measure computation were implemented using Gephi (v.0.8.2) [24]. The probability of reaching each state was also calculated using Markov chain simulations. Therefore, probabilistic Boolean networks (PBN) was used to specify more than one transition function per variable/component with specified probability values. It should be noted that the probabilities of all functions for each component sum up to one. The state transition graph was then reconstructed by choosing one function per component using probability values [22, 23]. In addition to all the eight versions of the present-day CD model, this analysis was also undertaken on the probabilistic version of the combined Boolean network (V1_V8) as shown in Table 3.

**Table 3:** The probabilistic version of the combined Boolean network of the present-day Central Dogma. The symbols “&”, “|” and “!” represent “AND”, “OR” and “NOT” operators.

| **Nodes** | **Logical rule** | **Probabilities** |
| --- | --- | --- |
| *Activator* | Protein \| Activator | 1 |
| *DegProtein* | Protein & !DegProtein | 1 |
| *DegRNA* | miRNA & Protein & !DegProtein | 1 |
| *DNA* | (miRNA & Protein) \| DNA | 0.1 |
|  | (!miRNA & Protein) \| DNA | 0.9 |
| *miRNA* | (DNA & Protein \| mRNA) & !DegRNA | 0.8 |
|  | (DNA & !Protein \| mRNA) & !DegRNA | 0.2 |
| *mRNA* | DNA & Protein & !DegRNA | 0.8 |
|  | DNA & !Protein & !DegRNA | 0.2 |
| *Protein* | (mRNA & Activator \| Protein) & !DegProtein | 1 |

Finally, the plausibility of the reconstructed models was assessed using two different measures of robustness to noise and mismeasurements. The normalized Hamming distance, which is the fraction of different Boolean bits between the states and the corresponding perturbed copies was obtained from 100 randomly generated copies [25]. The Gini index, an index of homogeneity in the in-degree property of the nodes, was calculated in the state-transition graph [26]. Both measures were statistically assessed based on the z-test and randomized version of the networks to demonstrate statistical significance of difference (Supplementary file 2 and 3).

## Results and discussion

### Synchronous modeling

We found by simulation that, with the synchronous update, the old conceptual model possesses 11 simple attractors (or fixed points) out of the 16 possible initial conditions (Table 4), indicating that 69% of all possible states were steady state forms, in which eight and seven fixed points were observed in the Activatory and Inhibitory models respectively. Four of these fixed points occurred in both the Activatory and Inhibitory gene expression versions (A, B, C and D). A null fixed point (state A) indicates a state where all the four nodes are in OFF state. This steady state (SS) was reachable in 12.5% (2 out of 16) of the initial states. The three other SS (B, C and D) did not include any transcribable genes, while mRNA, Protein and Activator remained unchanged. Among the four other fixed points (E, F, G and H) which reached only in the Activatory model, the G state had a larger basin of attraction (1110→1101→1111) and thus consistent with that expected from the 1965 model. Since mRNA and Protein turnover were not considered in this model, the G and H states were assumed as fixed points. The J state was another case showing consistency with the 1965 model expectation when Protein negatively regulates mRNA transcription. This state had the largest basin of attraction in the Inhibitory model (1100→1110→1111→1101). Analogous to states G and H, state I was also an artefact of ignoring mRNA turnover. No limit cycle was observed in these models based on the assumptions that were incorporated in the 1965 model. Furthermore, D, H and K states were specific to the 1965 CD model and do not fit with the current understanding of CD. For instance, it was not possible to reach any steady state in the attractor analysis of the present-day model if the state of Activator/DNA/mRNA was zero and Protein was ON similar to D.

**Table 4:** The attractors of the Boolean model as depicted in Fig. 1B obtained from the synchronous update method. In each version of the model, the number of single SSs and the corresponding normalized basin of attractions are demonstrated. The color gradient from red, yellow to green illustrate the lowest to the highest values, respectively. A to K attractors are encoded in the following order: Activator/DNA/mRNA/Protein. The D*, H* and K* denote specific states pertaining to this Central Dogma modeling.

| Central Dogma 1965 | Description | No. SSs with one state | A | B | C | D* | E | F | G | H* | I | J | K* |
| --- | --- | --- | --- | --- | --- | --- | --- | --- | --- | --- | --- | --- | --- |
|  |  |  | 0000 | 1000 | 1001 | 0001 | 0100 | 1100 | 1111 | 0111 | 0110 | 1101 | 0101 |
| V1 | Activatory gene expression | 8 | 12.5 | 6.25 | 18.75 | 12.5 | 12.5 | 6.25 | 18.75 | 12.5 | - | - | - |
| V2 | Inhibitory gene expression | 7 | 12.5 | 6.25 | 18.75 | 12.5 | - | - | - | - | 12.5 | 25 | 12.5 |
| Activator/DNA/mRNA/Protein |  |  |  |  |  |  |  |  |  |  |  |  |  |

Our results indicate that by including steady states in the present-day model of CD, obtained by the synchronous updating method, this model becomes more complex, hence more consistent with the current understanding of molecular biology. Overall, 12% (15 out of 128) of the possible initial states fell into the attractor set (Table 5). Compared with the 1965 model, which showed almost six-fold attractor percentage, the present-day model has a far more dynamic nature and thus a more realistic model. However, similar to the 1965 model, the null fixed point (state A) and the Activator ON state (state B) were observed in all eight versions with similar basins of attraction. There were five other fixed points that clustered the versions in the present-day model into three classes. The C and D states were observed in the Activatory (V1) and Gene silencing (V4) models which seems that without the basal expression of Protein, these models have similar behaviors with a final outcome of system collapse. State E clustered all versions of the model which contained inhibition of mRNA expression (i.e. mRNA expression inhibition (V2), Gene silencing & mRNA expression inhibition (V5), mRNA & miRNA expression inhibition (V7) and Inhibitory (V8) versions. Although the turnover of RNA species was considered in this model, the intactness of mRNA in the model is in line with amino acid paucity which is not included in the model. In addition, even if the translation had occurred and amino acids were in excess, the proteins were nonetheless not activated due to the absence of the Activator. Therefore, it is not surprising for state E to have appropriated a small fraction of all states (12.5%; 16 out of 128) in all the four mentioned versions. States F and G were analogous to state E but were restricted to the versions of the model which included inhibition of miRNA expression only (i.e. miRNA expression inhibition (V3) and gene silencing and miRNA expression inhibition (V6)).

**Table 5:** The attractors of the Boolean model depicted in Fig. 1C obtained from the synchronous update method. In each version of the model, demonstrated are the number of single SSs, SSs with multiple states (limit cycles) and corresponding normalized basin of attractions. The color gradient from red, yellow to green illustrate the lowest to highest values. The attractors A to I are encoded in the following order:

| Present-day Central Dogma | System name | No. of SS with one state | No. of SS with multiple states | A | B | C | D | E | F | G | H | I |
| --- | --- | --- | --- | --- | --- | --- | --- | --- | --- | --- | --- | --- |
|  |  |  |  | 0000000 | 1000000 | 0001000 | 1001000 | 0001110 | 0001100 | 1001100 | 1001000<br>><br>1001010<br>><br>1001111<br>><br>1111101 | 1001000<br>><br>1001110<br>><br>1001111<br>><br>1111101 |
| V1 | Activatory | 4 | 0 | 12.5 | 21.88 | 12.5 | 53.12 | - | - | - | - | - |
| V2 | mRNA expression inhibition | 3 | 1 | 12.5 | 21.88 | - | - | 12.5 | - | - | 53.12 | - |
| V3 | miRNA expression inhibition | 4 | 0 | 12.5 | 21.88 | - | - | - | 12.5 | 53.12 | - | - |
| V4 | Gene silencing | 4 | 0 | 12.5 | 17.19 | 12.5 | 57.81 | - | - | - | - | - |
| V5 | Gene silencing & mRNA expression inhibition | 3 | 1 | 12.5 | 17.19 | - | - | 12.5 | - | - | 57.81 | - |
| V6 | Gene silencing & miRNA expression inhibition | 4 | 0 | 12.5 | 17.19 | - | - | - | 12.5 | 57.81 | - | - |
| V7 | miRNA & mRNA expression inhibition | 3 | 1 | 12.5 | 21.88 | - | - | 12.5 | - | - | - | 53.12 |
| V8 | Inhibitory | 3 | 1 | 12.5 | 17.19 | - | - | 12.5 | - | - | - | 57.81 |
| Activator/DegProtein/DegRNA/DNA/miRNA/mRNA/Protein |  |  |  |  |  |  |  |  |  |  |  |  |

Unlike the previous models, there were two limit cycles playing the attractor role in this simulation. State H is the common cycle between mRNA expression inhibition (V2) and Gene silencing & mRNA expression inhibition (V5) attracting more than 50% of all initial states. Similarly, state I is the common cycle between those having inhibition of RNA species expression namely mRNA & miRNA expression inhibition (V7) and Inhibitory (V8). A common trait in all limit cycles was the constant presence of DNA and Activator. It is, therefore, expected that steady states are related to the long half-life components of the biological systems.

### Asynchronous modeling

Using the asynchronous Boolean dynamic approach, we analyzed the present-day model of CD to explore its attractors. Tantamount to the fixed points found in the synchronous updating model, there were seven simple attractors (states A to G) with profiles identical to previous attractors (Table 6). However, the complex (loose) attractor was found to be different from the limit cycle attractors of the synchronous approach and appeared to be more convoluted. This large complex attractor comprised 32 nodes and 84 edges (state H) (see Table 6 and Fig. 2) where nodes played various roles in the graph. Given that the node size is proportional to betweenness centrality, larger nodes (i.e. states) are those with more presence in transitions from one state to another. With respect to closeness centrality (represented by node color), darker nodes are those with a higher closeness centrality measure and thus more accessible or close to other nodes. The node 1111111, which is the full ON state of all components, had the highest closeness in this loose attractor, however, not the highest betweenness. The highest betweenness value was observed for the state in which the degradation components were not active (i.e. 1001111). Furthermore, simulation data suggest that the inhibition of mRNA expression alone or in combination with miRNA expression inhibition and gene silencing is the main contributor to oscillatory behavior of the molecular machinery.

**Table 6:** The attractors of the Boolean model depicted in Fig. 1C obtained from the asynchronous updating method. Each version of the model illustrated the presence of simple attractors, the complex (loose) attractor and corresponding edge numbers in the latter.

| Present-day<br>Central<br>Dogma | System name | Simple attractor |  |  |  |  |  |  | Complex (loose) attractor |
| --- | --- | --- | --- | --- | --- | --- | --- | --- | --- |
|  |  | A | B | C | D | E | F | G | H |
|  |  | 0000000 | 1000000 | 0001000 | 1001000 | 0001110 | 1001100 | 0001100 | 1111111,1011111,1101111,1001111,1111011,1011011,1101011,1001011,1111101,1011101,1101101,1001101,1111001,1011001,1101001,1001001,1111110,1011110,1101110,1001110,1111010,1011010,1101010,1001010,1111100,1011100,1101100,1001100,1111000,1011000,1101000,1001000 |
| V1 | Activatory | ✓ | ✓ | ✓ | ✓ |  |  |  |  |
| V2 | mRNA expression inhibition | ✓ | ✓ |  |  | ✓ |  |  | 32 |
| V3 | miRNA expression inhibition | ✓ | ✓ |  |  |  | ✓ | ✓ |  |
| V4 | Gene silencing | ✓ | ✓ | ✓ | ✓ |  |  |  |  |
| V5 | Gene silencing & mRNA expression inhibition | ✓ | ✓ |  |  | ✓ |  |  | 32 |
| V6 | Gene silencing & miRNA expression inhibition | ✓ | ✓ |  |  |  | ✓ | ✓ |  |
| V7 | miRNA & mRNA expression inhibition | ✓ | ✓ |  |  | ✓ |  |  | 32 |
| V8 | Inhibitory | ✓ | ✓ |  |  | ✓ |  |  | 32 |
| Activator/DegProtein/DegRNA/DNA/miRNA/mRNA/Protein |  |  |  |  |  |  |  |  |  |

**Fig. 2.**
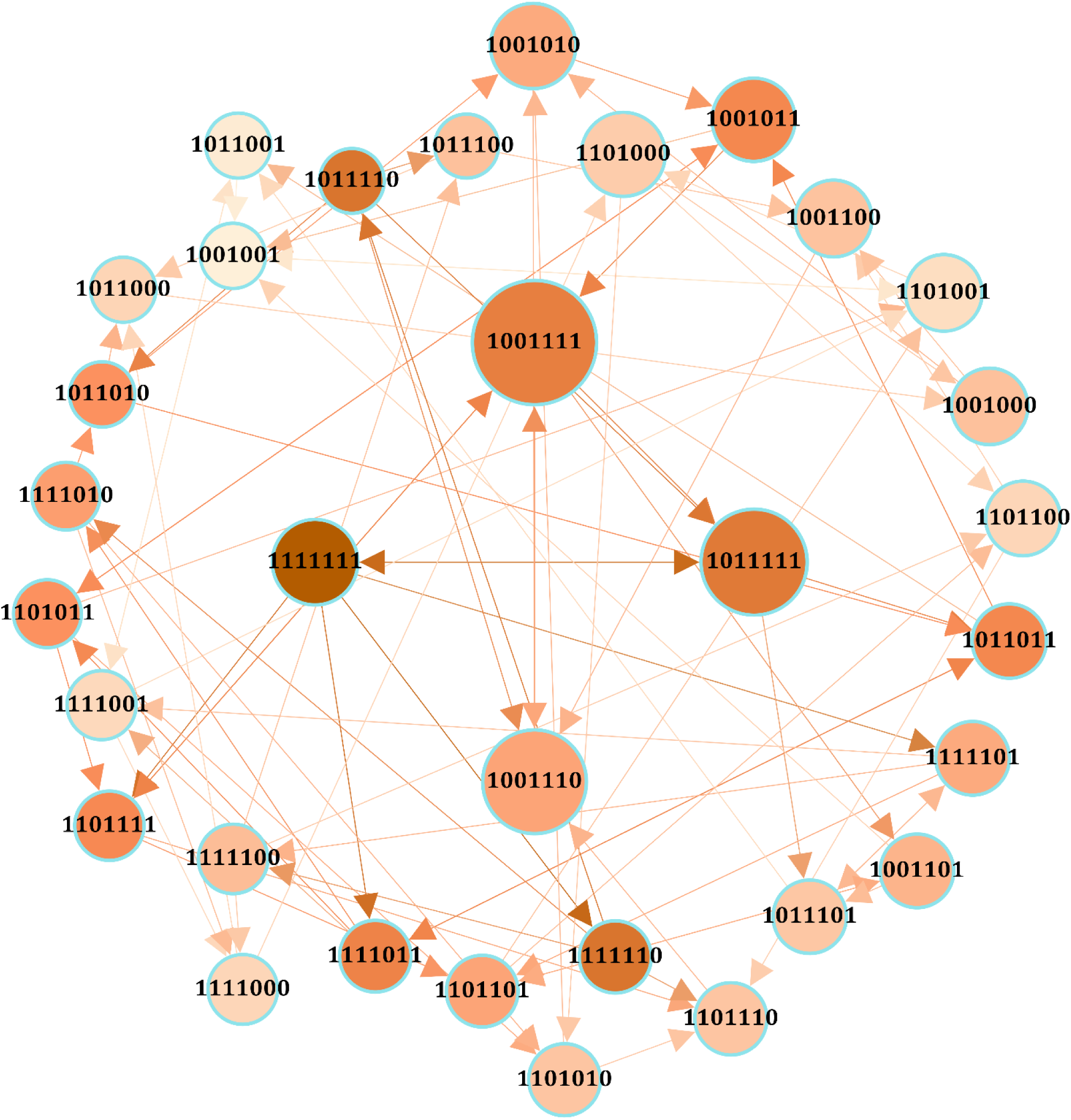
Network visualization of the complex (loose) attractor in the present-day model. Node size and color are proportional to betweenness and closeness centrality measures respectively.

To estimate the probability of reaching an attractor following a large number of iterations, we performed a Markov chain simulation on the present-day CD model. In Table 7, the states reached at the end of the simulation (attractors) are listed for all versions of the model. In addition, the combined version (V1_V8) was added to the analysis to systematically study the global dynamic network by incorporating all versions, an approach which is robust in tackling uncertainty. In all versions, it was equally probable to reach the null fixed point and fixed point with only the Activator ON. Based on the activation/inhibition functions in each version, other states occurred with different probabilities. Apart from these two states (0000000, 1000000), as expected, 1001000 had the highest probability in the Activatory (V1), Gene silencing (V4) and combined V1_V8 versions The states 1111101 and 1001100 were the most probable states to reach in the inhibited mRNA expression versions (V2 and V5) and the inhibited miRNA expression versions (V3 and V6) respectively. The 1111101 was, as expected, the most probable state for V7 and V8 versions of the model. A number of states were unique to the combined version (V1_V8), including 0001010, 1001101, 1111001, 1111011 and 1111111. This demonstrates that combining independent versions of model results in observing undetected states in single versions.

**Table 7:** The absorption probabilities for the Markov chain corresponding to the Boolean model depicted in Fig. 1C in all versions. These probabilities were estimated based on 1000 iterations. The last column represents the absorption probabilities of the probabilistic Boolean network of the present-day Central Dogma (See table 2).

| Activator/DegProtein/<br>DegRNA/DNA/miRNA/<br>mRNA/Protein | V1 | V2 | V3 | V4 | V5 | V6 | V7 | V8 | Mixture<br>V1_V8 |
| --- | --- | --- | --- | --- | --- | --- | --- | --- | --- |
| 0000000 | 0.125 | 0.125 | 0.125 | 0.125 | 0.125 | 0.125 | 0.125 | 0.125 | 0.125 |
| 0001000 | 0.125 |  |  | 0.125 |  |  |  |  | 0.064 |
| 0001010 |  |  |  |  |  |  |  |  | <u>0.016</u> |
| 0001100 |  |  | 0.125 |  |  | 0.125 |  |  | 0.036 |
| 0001110 |  | 0.125 |  |  | 0.125 |  | 0.125 | 0.125 | 0.009 |
| 1000000 | 0.219 | 0.219 | 0.219 | 0.172 | 0.172 | 0.172 | 0.219 | 0.172 | 0.177 |
| 1001000 | 0.531 | 0.047 |  | 0.578 | 0.047 |  | 0.047 | 0.047 | 0.301 |
| 1001010 |  | 0.047 |  |  | 0.063 |  |  |  | 0.057 |
| 1001100 |  |  | 0.531 |  |  | 0.578 |  |  | 0.057 |
| 1001101 |  |  |  |  |  |  |  |  | <u>0.057</u> |
| 1001110 |  |  |  |  |  |  | 0.047 | 0.063 | 0.014 |
| 1001111 |  | 0.203 |  |  | 0.234 |  | 0.203 | 0.234 | 0.014 |
| 1111001 |  |  |  |  |  |  |  |  | <u>0.002</u> |
| 1111011 |  |  |  |  |  |  |  |  | <u>0.009</u> |
| 1111101 |  | 0.234 |  |  | 0.234 |  | 0.234 | 0.234 | 0.012 |
| 1111111 |  |  |  |  |  |  |  |  | <u>0.048</u> |

### Robustness of the reconstructed model

To assess the plausibility of the CD models, we tested model robustness to noise and mismeasurements in all versions of the 1965 and present-day models of CD. As shown in Table 8, the present-day model was more robust than the previous model. In specifc, the Hamming distance was significantly smaller in all eight versions of this model, meaning that by applying noise to the network states in these models, attractors remained largely unperturbed. The measure of in-degree homogeneity (Gini index) was assessed in the state transition graph of all versions in both models and was further compared with the randomized graph to assess statistical significance. Generally, as expected in biological networks, the present-day model is enriched with many low in-degree nodes along with a few high in-degree nodes in the state transition graph, indicating that the updated model is more consistent with the reality of biological processes.

**Table 8:** The robustness analysis of the reconstructed models of the Central Dogma.

| Model Versions | Hamming distance | P-value | Gini Index | P-value of the Gini Index |
| --- | --- | --- | --- | --- |
| **1965** |  |  |  |  |
| V1 | 0.271 | 0.19 | 0.588 | 0.52 |
| V2 | 0.275 | 0.1 | 0.575 | 0.61 |
| **2009** |  |  |  |  |
| V1 | 0.162 | <0.001 | 0.918 | <0.001 |
| V2 | 0.149 | <0.001 | 0.908 | <0.001 |
| V3 | 0.151 | <0.001 | 0.904 | <0.001 |
| V4 | 0.151 | 0.01 | 0.918 | <0.001 |
| V5 | 0.144 | 0.01 | 0.904 | <0.001 |
| V6 | 0.148 | <0.001 | 0.908 | <0.001 |
| V7 | 0.146 | <0.001 | 0.921 | <0.001 |
| V8 | 0.145 | <0.001 | 0.921 | <0.001 |

## Conclusion

Biology, in this contemporary era, is an edifice made up of two primary axioms as building blocks, Darwin’s theory of evolution and the Central Dogma of information flow in molecular biology [3]. Although the overall structure of the latter (i.e. information flow from DNA-> RNA-> Protein) has remained intact over the past half-century, the complexity of this cascade has surprisingly increased. Discovering diverse species of RNA with distinguishing roles, as well as dynamic features of DNA, mRNA and proteins, from cell to cell and over time, contribute to this complexity [4, 27, 28]. The Boolean network modeling, as a dynamic framework, was essential to the development of the CD models and providing further insight into gene regulation. We show that in addition to predicting reliable steady states, the present-day model of CD is more robust against and insensitive to noise than the 1965 model. Using this mathematical modeling approach, we put forward a logic-based expression of our conceptual view of molecular biology [29], thus, not only validating our abstract understanding of molecular biology, but it also improves our predictive power of biological processes.

## Supplementary file

**Supplementary file 1: Detailed description of functions used in the developed CD models.**

**Supplementary file 2: The R script containing all computational codes used in the present study.**

**Supplementary file 3: The SBML files pertaining to different Boolean networks reconstructed and simulated in this study.**

